# Spatiotemporal analysis of host cell modification during herpes simplex virus 1 replication

**DOI:** 10.1101/2019.12.22.872002

**Authors:** Katharina M. Scherer, James D. Manton, Timothy K. Soh, Luca Mascheroni, Viv Connor, Colin M. Crump, Clemens F. Kaminski

## Abstract

Herpesviruses are large and complex viruses, and have a long evolutionary history with their host species. One poorly understood, but important, area in virus-host interaction is the extensive remodelling of intracellular organelles and morphology that occurs during active viral replication. We have constructed a recombinant reporter virus that allowed us to classify four distinct stages in the infection cycle of herpes simplex virus 1. Single-cell analysis of the impact of the viral load on replication dynamics demonstrates the utility of this times-tamping method for tracking the infection cycle. High-resolution microscopy analysis of cellular structures in live and fixed cells in concert with our dual-fluorescent reporter virus have enabled us to generate a detailed overview over the spatial and temporal organisation of the cytoskeleton and organelles during viral replication. This provides the first systems-level analysis of the morphological changes involved in viral replication.

## Introduction

Herpesviruses are large and complex DNA viruses which are typified by their ability to establish both lytic and latent infection cycles. For the prototypical herpesvirus, herpes simplex virus 1 (HSV-1), lytic infections occur in epithelial cells whereas life-long latent infection is established in sensory neurons. Herpesviruses are among the most complex viruses, with a large repertoire of genes that allows for an intricate relationship with their host. During the lytic infection cycle, the activity of herpesvirus gene products profoundly alters cellular physiology, causing dramatic modifications of the infected cell that convert it into an efficient virus-producing factory. These changes to the intracellular environment arise as a consequence of the manipulation of a range of cellular functions and are intimately linked to key processes in the viral replication cycle such as virus assembly and egress [1–3]. However, there is relatively little understanding of the dynamics of morphological changes of infected cells and how they relate to distinct phases of the herpesvirus replication cycle [4].

There are a few examples in the literature that show how the biophysical analysis of cellular remodeling events has led to a better understanding of the mechanisms of HSV-1 replication. The best studied example is the nucleus. During replication, HSV-1 remodels the nuclear architecture by increasing the volume of the nucleus [5] and rearranging the host chromatin to facilitate egress of capsids [6, 7]. The impact of HSV-1 infection on the cytoskeleton has also been studied previously, with stabilisation of microtubule subsets and dramatic reorganisation of the microtubule network having been described at late stages of infection [8]. Membrane compartments are affected during HSV-1 infection as well. Examples include the concentration of the endoplasmic reticulum (ER) around the nucleus [9], the alteration of mitochondrial dynamics [10], and the fragmentation of the Golgi apparatus [11].

However, consistent and systematic data is lacking for this important area in host-pathogen interaction. The focus on usually one or only a few structures as well as the use of different experimental conditions and techniques renders comparison and correlation between results from previous studies nearly impossible. Furthermore, investigation of the dynamic aspects of morphological alterations over the whole course of the herpesvirus replication cycle has been neglected. We sought to address these gaps in knowledge and provide a comprehensive understanding of the spatiotemporal changes in cell morphology that are induced during lytic herpesvirus infection.

There are several challenges which have limited previous studies. First, the asynchronous progression of infection yields large variation between single cells, and even with a high multiplicity of infection individual cells reside in different phases in the replication cycle. Without a direct readout for the stage of infection at the single cell level, correlating observed changes in cellular morphology with phases of viral replication is fraught with difficulty and prone to misinterpretation. Furthermore, it has become apparent over the last few years that intracellular membrane compartments engage in extensive communication, often through direct contact sites (described as the organelle interactome [12]). This new perspective on inter-organelle communication will influence our understanding of herpesvirus morphogenesis, but a systematic approach to map organelle changes over the time course of infection is missing. Finally, while imaging is the ideal tool to study intracellular cytoskeletal and organelle morphology, it is technically challenging to capture cellular rearrangements that take place over the whole viral replication cycle (12 hours for HSV-1) in great detail. Either a multiplexed acquisition with multiple color channels (as shown by Valm et al. [12]) has to be used together with time-lapse imaging over several hours, or a universal reference point or temporal marker would be needed that allows correlating different imaging sets.

In this study, we have characterised the spatiotemporal morphology changes for a broad range of organelles and the cytoskeleton on a single cell level in relation to the replication cycle of HSV-1 using state-of-the-art fluorescence microscopy techniques. We have constructed a dual-fluorescent reporter virus that expresses eYFP-fused to the immediate early protein ICP0 and mCherry-fused to glycoprotein C (gC), a true late protein. The distinct temporal and spatial expression patterns of these two fluorescently tagged reporter proteins provide an intrinsic timestamp enabling simple classification of four clear stages of infection. This novel classification based on visual expression patterns is ideally suited to describe the spatiotemporal remodeling processes. Combining this approach with structured illumination microscopy (SIM) and expansion microscopy has enabled us to map virus-induced structural re-arrangement of the cytoskeleton, mitochondria, and secretory pathway compartments in great detail. We have uncovered a concerted and dramatic relocation of early endosomes, mitochondria and microtubules during transition from early to late gene expression, which suggests that the reshaping of these compartments is required for virus assembly. We have also discovered that ICP0 colocalises with the microtubule organising centre (MTOC) during early stages of infection, suggesting a central role for this viral enzyme in microtubule reorganisation.

## Results

### Timestamping the viral replication state

HSV-1 replication is a dynamic process that spans a period of 6–12 hours from the entry of virions into the host cell until egress of newly formed virions. A measure that is usually used to indicate the state of replication during an experiment is the hours post infection (hpi). However, experimental conditions such as the multiplicity of infection (MOI), as well as the cell type used, influence the dynamics of replication. In addition, the asynchronous nature of infection in a cell population makes it impossible to determine the exact viral replication state within an individual cell just by indicating the hpi. To address these issues, we developed a fluorescent reporter virus for a direct visual readout of the replication state on the single cell level (Figure 1).

**Figure 1:**
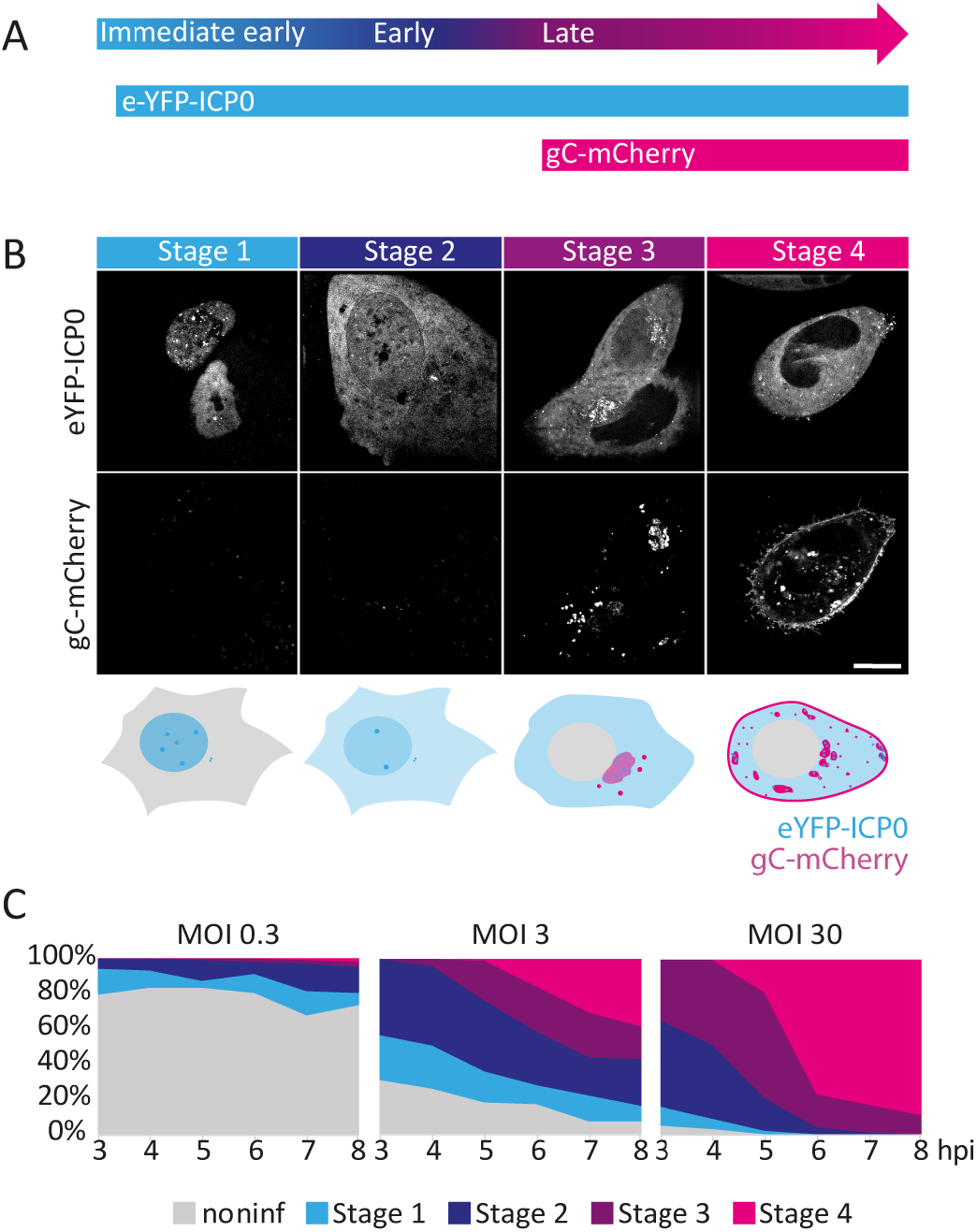
Timestamping the viral replication state inside individual cells. **A** Schematic illustration of the engineered reporter virus expression pattern. Three phases of the viral transcription cascade — immediate early, early and late — are characterised by the temporal expression of HSV-1 genes during lytic infection. **B** Structured illumination microscopy (SIM) images of HFF cells infected with eYFP-ICP0/gC-mCherry HSV-1. The cells were imaged at 3.5, 5.5, 7.5 and 9.5 hours post infection (hpi). Appearance and localisation of the two fluorescently labelled viral proteins can be correlated with four distinct stages in infection (stage 1 – stage 4) as depicted in the illustration below the micrographs. Scale bar 10 μm. **C** Progression of infection was measured on the single cell level for different multiplicities of infection (MOIs) by classification of > 150 cells per time point into the four stages of infection for 3–8 hpi.

As an indicator of early stages of infection we fluorescently labeled ICP0 with eYFP. ICP0 is one of five immediate early proteins in HSV-1, and it begins to be expressed as soon as the viral genome is deposited inside the nucleus of the host cell. In addition, glycoprotein C (gC), a true late protein, was labeled with mCherry to serve as a marker for the late stages of HSV-1 replication. We used time-lapse imaging to confirm the functionality of our construct (Supplementary Video 1). As expected, at 2–3 hpi cells start to exhibit fluorescence from eYFP-ICP0. At first, ICP0 is located in the nucleus, but translocates to the cytoplasm with further progression of the infection[13]. At later stages of infection, additional fluorescence signal from gC-mCherry appears. As a virion envelope protein, gC localises to sites of secondary envelopment. We observe that it is first located in a perinuclear membrane organelle before spreading into more dispersed puncta in the cytoplasm and to the plasma membrane late in infection.

In order to map the appearance and re-distribution of both proteins over the course of infection with high detail and contrast, we performed structured illumination microscopy (SIM). By doing so, we identified characteristic features that appear in a time-dependent pattern during HSV-1 replication. We used these visual features to define criteria by which cells could be sorted into four different categories which correspond to distinct stages in the HSV-1 replication cycle. Representative images are shown in (Figure 1B). More examples are displayed in Supplementary Figure 1.

During stage 1, ICP0 is located almost exclusively in the nucleus, and localises to discrete nuclear structures reminiscent of nuclear domain 10 (ND10) bodies. ICP0 disrupts ND10 body function by causing degradation of components such as PML and Sp100 [14]. The second stage in infection is characterised by translocation of IPC0 from the nucleus to the cytoplasm. This stage can be visually distinguished from stage 1 by an increase in the cytoplasmic eYFP signal above the background level and simultaneous decrease of the nuclear fluorescence signal. Supplementary Figure 1 shows several examples of snapshots taken of cells during stage 2 with varying ratios of nuclear-to-cytoplasmic eYFP-ICP0 intensities. During stage 3, when transition of ICP0 to the cytoplasm is completed, mCherry fluorescence signal appears due to expression of gC. gC is first localised at the perinuclear region, where ICP0 forms several small domains. As ICP0 and gC are virion components, these structures may represent compartments of virus assembly. Usually, cell morphology changes during stage 3, and cells start to round up. Stage 4 is marked by a spread of gC-enriched compartments in the cytoplasm and a high gC concentration at the plasma membrane. Cells at that stage are rounded up, show membrane blebbing as well as formation of thin membrane protrusions, and nuclei often exhibit indentations.

This new classification scheme was used to characterise the progression of infection on the single cell level for different multiplicities of infection (MOIs) (Figure 1C). The probability that a cell will become infected can be calculated using a Poisson distribution [15]. As the MOI increases, the percentages of cells infected with at least one viral particle also increases and amount to 26%, 95% and 100% for MOIs 0.3, 3 and 30, respectively. As Figure 1C shows, these predictions are consistent with the experimental findings of our singlecell assay. For all three MOIs, we observe at every hpi a combination of individual stages of infection. The plots furthermore show that progression of infection is faster for higher MOIs whereas for the low MOI (0.3) the number of noninfected cells remains constant over the observed period of hpi. Also, for the low MOI, the early stages in infection (stage 1 and 2) dominate until 5 hpi whereas for the high MOI 30, from 5 hpi on late stages in infection (stage 3 and 4) prevail. For the middle MOI 3, which was used throughout all further experiments in this study, we find a mixture of all stages from 5 hpi on.

### Re-organisation of the cytoskeleton

The cytoskeleton plays an important role for the replication of HSV-1. For effective viral assembly and egress, which take place during the late phases of replication, HSV-1 uses microtubules to transport nucleocapsids to the sites of secondary envelopment - membrane compartments likely originating from endosomes and the trans-Golgi network (TGN) (see reviews by Johnson and Baines [16] and Owens et al. [17]). However, it is also known that the cytoskeletal architecture is extensively remodelled during HSV-1 replication [18, 19]. By use of our ‘timestamp’ reporter virus, we investigated when and in which way microtubule architecture changed during replication. For imaging, microtubules were stained with SiR-tubulin in cells infected with the reporter virus. SiR-tubulin consists of the fluorophore silicon rhodamine (SiR) conjugated to the microtubule binding drug Docetaxel. It is a marker for all types of tubulin including microtubule filaments (consisting of alpha- and beta-tubulin) and the centrosome/microtubule organising center (MTOC, consisting of gamma-tubulin). Figure 2A shows the re-modelling of the microtubule network in representative images for the different stages of infection. At the second stage, microtubules start to disconnect from the MTOC. During transition from stage 2 to stage 3, microtubules are cleared from the perinuclear region and start to form thick bundles around the perinuclear region. Furthermore, SiR-tubulin staining of the MTOC can no longer be observed at stage 4. Interestingly, we observed that ICP0 colocalises with the MTOC during the early replication stages (Figure 2B). The direct recruitment of ICP0 to this organelle may explain the observed loss of the MTOC and the published role of ICP0 in modifying the microtubule network in HSV-1 infected cells [20].

**Figure 2:**
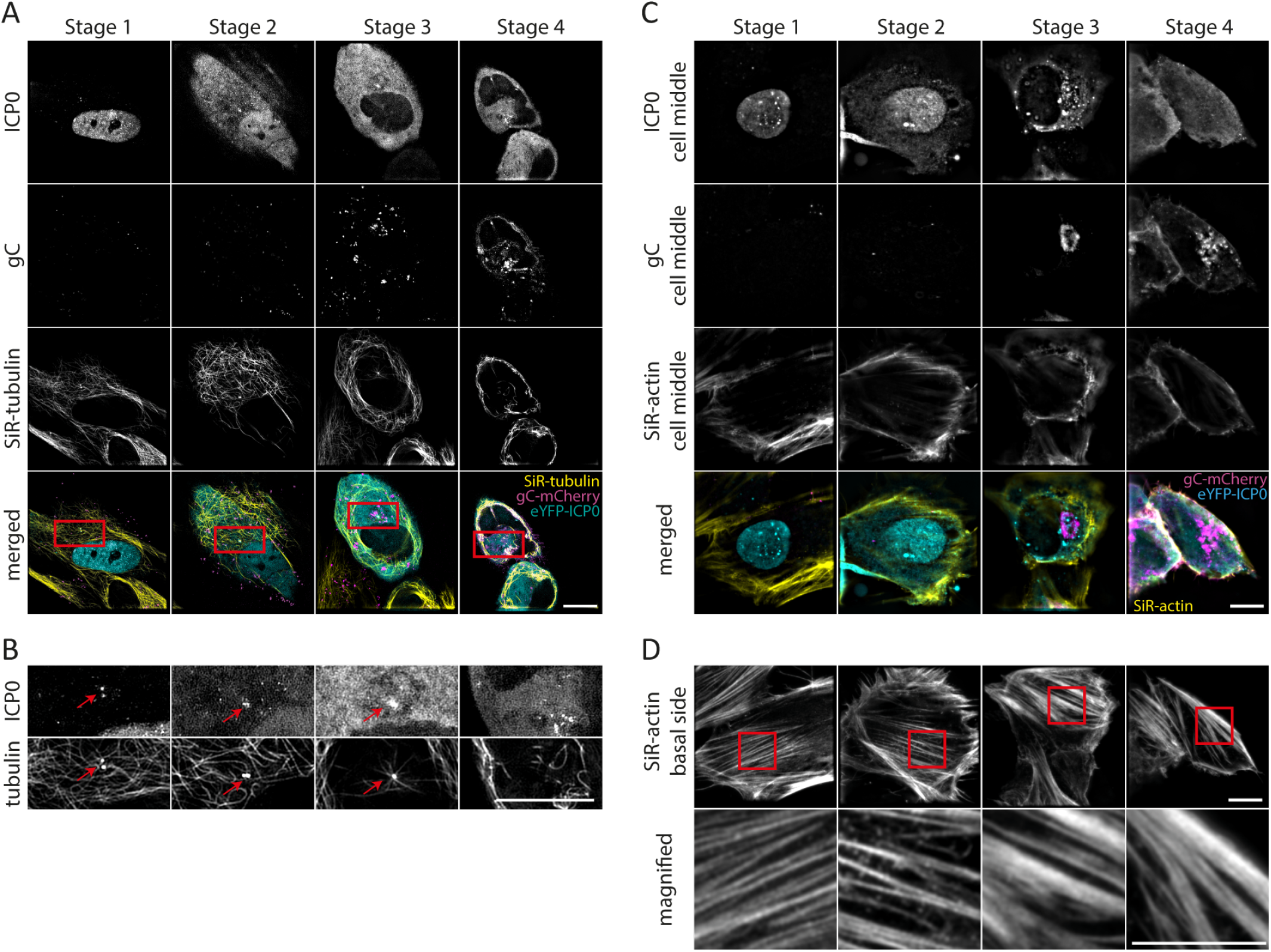
Morphological changes of cytoskeletal components dependent on the stage of infection. HFF cells were infected with eYFP-ICP0/gC-mCherry HSV-1, stained with SiR-tubulin (**A, B**) or SiR-actin (**C,D**), respectively, and imaged at a custom-built SIM microscope. **A** The two upper rows show the two viral proteins ICP0 and gC. Images were used to classify the stage of infection in individual cells. Third row shows the re-organisation of microtubules during infection. At late stages in infection, microtubules are bundled and excluded from the perinuclear region. Bottom row shows merged images of all three channels. **B** Enlarged view of the perinuclear region as marked with red boxes in (**A**) (top row: ICP0, bottom row: SiR-tubulin). Arrows highlight the colocalisation of ICP0 with the microtubule organising center (MTOC) at the centrosome during the early phase in replication. **C** For classification purposes, three-colour images of ICP0, gC and SiR-actin were taken at a middle section of each cell. Bottom row shows merged images of all three channels. **D** Actin stress fibers undergo remodelling during infection. Bottom row shows enlarged view of actin stress fibers (areas marked with red boxes in upper row). Scale bar 10 μm.

Actin is an important structural component of the cell, and also important for intracellular transport. Since several studies have reported that herpesvirus exploits actin and actin-associated myosin motors for viral entry, intranuclear transport of capsids, and egress, we were interested in any morphological changes that HSV-1 induces to the actin cortex and the time at which they occur during replication. We first imaged the middle section of infected cells stained with SiR-actin to classify the stage of infection (Figure 2C). By means of the actin stain the cell outline is clearly identifiable, and shows that cells are rounded up during late stages in infection (stage 3 and 4). This morphological change is reflected in the reorganisation of the stress fibers in the actin cortex at the cell base. We observe a transition from thin stress fibers early in infection (stage 1) to thick fibers (stage 4) (Figure 2D). The morphological changes seem to be initiated during stage 2 where many of the stress fibers become branched.

### Transport pathways and potential envelopment compartments

Next, we studied the infection-induced re-organisation of compartments involved in glycoprotein trafficking and virus assembly. The assembly of herpesviruses is known to involve the wrapping/budding of nucleocapsids, together with the complex layer of tegument proteins, at membranes derived from post-Golgi endocytic compartments. Markers of the Golgi apparatus and early endosomes were labeled by immunostaining (58K protein for the Golgi and EEA1 for early endosomes) or tagged with the autofluorescent protein mIFP which is compatible with the eYFP and mCherry tags of ICP0 and gC (B4GAL-T1 for the Golgi apparatus and Rab4a for early endosomes).

The Golgi apparatus, located next to the nucleus and centrosome, is a key component of the secretory pathway. In HSV-1 infection, viral glycoproteins accumulate in the Golgi apparatus where they are modified before transport to the plasma membrane and endosomes. Immunostaining of the 58K Golgi protein yielded a strong fluorescence signal which allowed the clear identification of the Golgi apparatus (Figure 3A). The mIFP-tagged B4G4l-T1 is concentrated at the juxtanuclear Golgi compartment, but is also found in vesicular structures throughout the cytoplasm (Supplementary Figure 2). The most apparent change in Golgi morphology takes place during very late stages in infection. At stage 4, the Golgi apparatus fragments into several smaller compartments and a great number of small structures, an event we observed with both Golgi markers (Figure 3A and Supplementary Figure 2). The dispersion of the Golgi complex also influences the distribution of gC. During stage 3, gC is mainly located at the Golgi compartment. The fragmentation of the Golgi apparatus leads to a spread of gC-enriched membrane compartments in the cytoplasm of the host cell. Interestingly, by measuring the area occupied by the Golgi apparatus prior to fragmentation at stage 4, we observed a continuous compaction of the compartment from stage 1 to stage 3 (Figure 3B). A change in shape can also be observed with the mIFP-tagged marker B4G4L-T1 where the Golgi complex appears to be more compact across stages 2 to 3.

**Figure 3:**
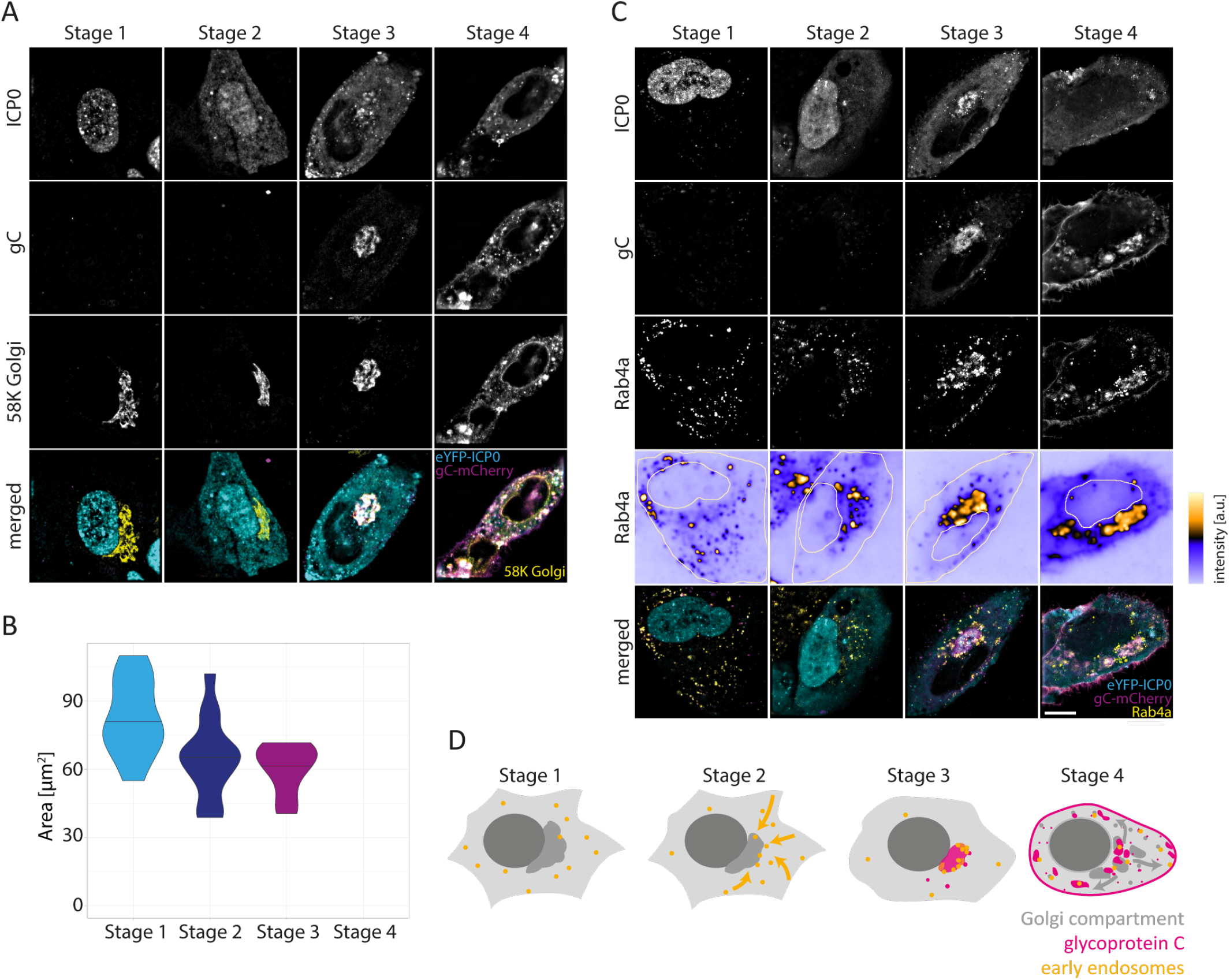
Remodelling of membrane compartments involved in glycoprotein trafficking and HSV-1 assembly. **A** HFF cells were infected with eYFP-ICP0/gC-mCherry HSV-1, fixed at 3, 6 and 9 hpi, and the Golgi apparatus was stained with an anti-58K Golgi protein antibody following standard immunofluorescence protocol. **B** The 58K Golgi protein marker was used to measure the size of the Golgi compartment over the time course of the replication cycle. The area occupied by the Golgi apparatus was calculated for at least 10 cells for each stage (stage 1: *n* = 11, stage 2: *n* = 14, stage 3: *n* = 11). The area continuously decreases from stage 1 to stage 3. During stage 4, the Golgi apparatus was fragmented in all imaged cells (n = 12) and no area was calculated. **C** Accumulation of early endosomes at the perinuclear region and spread correlated to fragmentation of perinuclear compartment. At stage 1, early endosomes are widely spread in the cytoplasm, but start to accumulate at the perinuclear region during stage 2. At stage 3, early endosomes are further concentrated at the gC-enriched perinuclear region, and Rab4a is found within the assembly compartment. Fragmentation of the assembly compartment leads to re-spread of early endosomes in the cytoplasm during stage 4. Scale bar 10 μm. **D** Schematic illustration of early endosome and assembly compartment reorganisation in relation to the Golgi complex.

The observed Golgi fragmentation is accompanied by a loss of the centrosome/MTOC (Figure 4A). To specifically label the centrosome, cells were immunostained for pericentrin. This organelle can clearly be observed during stages 1 to 3. However, it disappears during stage 4, similar to SiR-tubulin labeling of the MTOC. The loss of the centrosome due to Golgi fragmentation occurs together with the dispersal of gC within the cytoplasm and the plasma membrane due to Golgi fragmentation. Figure 4B shows an enlarged view of the perinuclear region. Consistent with Figure 2B we find that ICP0 localises to the centrosome during early stages in infection.

**Figure 4:**
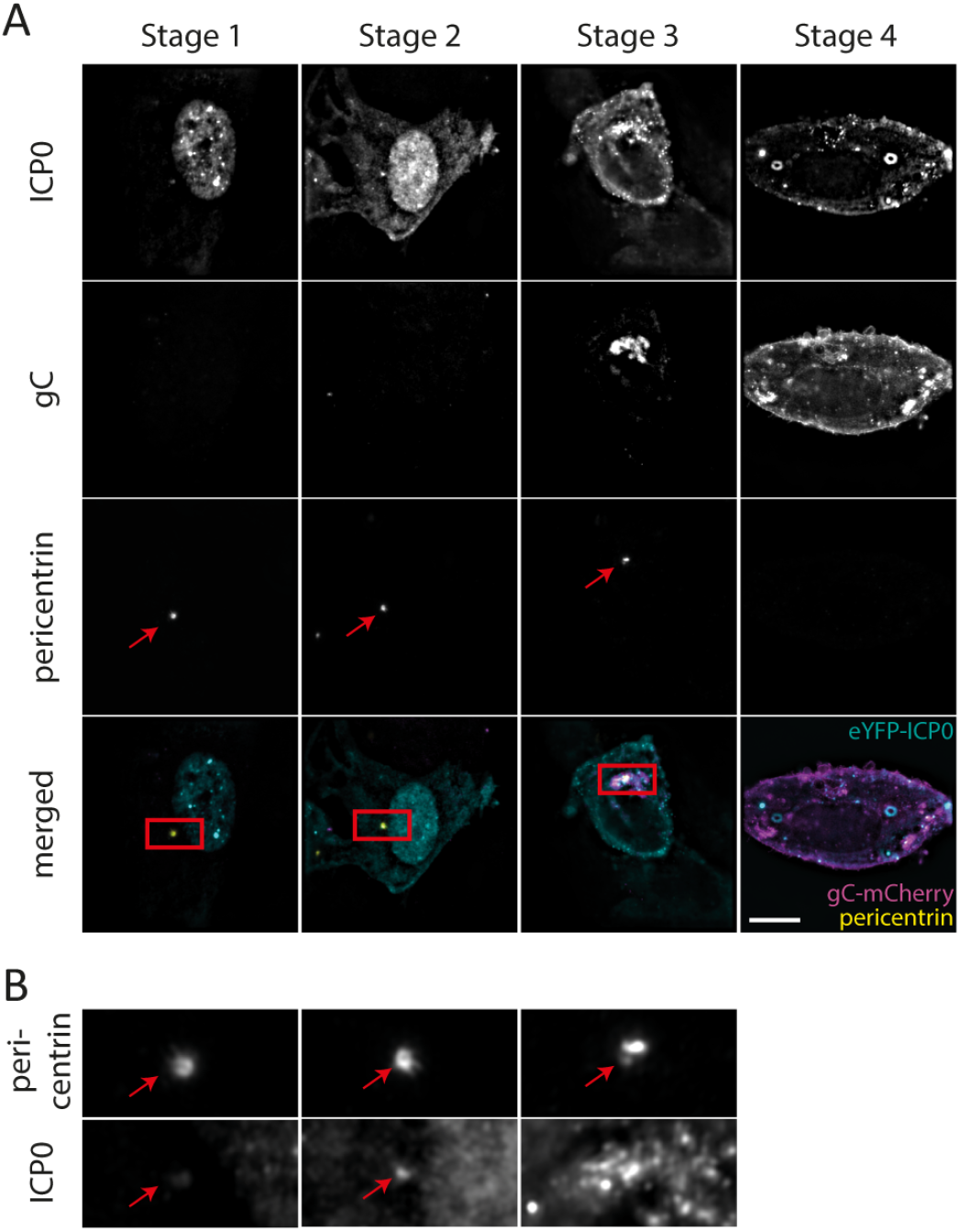
Disappearance of centrosome correlated to fragmentation of the Golgi apparatus. **A** HFF cells were infected with eYFP-ICP0/gC-mCherry HSV-1, fixed at 3.5, 5.5, 7.5 and 9.5 hpi, and the centrosome labeled with an antibody for pericentrin. **B** During stage 1 and 2, colocalisation of ICP0 with the centrosome can be detected. Scale bar 10 μm.

As potential sources of membrane for secondary envelopment [21, 22], endosomes likely play a key role in HSV-1 assembly. Therefore, understanding the spatial and temporal reorganisation of early endosomes during the replication cycle could help to further identify the mechanisms underlying HSV-1 assembly. In order to monitor the spatial distribution of early endosomes during the HSV-1 replication cycle, we generated a stable cell line expressing Rab4a tagged with the autofluorescent protein mIFP. Figure 3C shows that Rab4a localisation changes dramatically over the viral replication cycle. Rab4a vesicles are of small size, and characterised by a strong fluorescence signal. They relocate to the perinuclear region during an early stage in infection (stage 2). During stage 3, Rab4a vesicles are highly concentrated at and around the gC-enriched assembly compartment at the perinuclear region. Very late in infection at stage 4, Rab4a vesicles are redistributed within the cytoplasm. However, we also detect subregions of weaker Rab4a fluorescence signals within regions that become or are enriched with gC. Starting at stage 2, the effect is more distinct for stages 3 and 4 where a diffuse Rab4a signal is found at the gC-rich perinuclear and later fragmented compartments. A very similar behavior at all stages was also observed for a second early endosome marker, EEA1, which was monitored in fixed cells using immunostaining (Supplementary Figure 3). These observations suggest that early endosomes partially fuse with other membranes near the centrosome, perhaps indicating formation of secondary envelopment compartments.

We also studied the location of lysosomes during infection. Lysosomes are trafficked between the perinuclear region and cell periphery via microtubule-driven transport, and their position is influenced, for example, by nutrient starvation. Imaging of lysosomes stained with SiR-lysosome in live cells revealed a similar behavior to early endosomes (Supplementary Figure 4). Lysosomes are directed towards the perinuclear region during early stage in infection, and accumulate near gC-rich membrane compartments late in infection. This suggests concentration of all endocytic compartments at the perinuclear assembly compartment at stage 3, followed by dispersal during stage 4.

### Antiviral and inflammatory signalling platforms

A link between mitochondrial dynamics and viral infections has been reported for a wide range of viruses (see review [23]). Mitochondria also act as platforms for antiviral immunity through activation of the retinoic acid-inducible gene I (RIG-I)-like receptors signal transduction pathway [24]. To observe changes in mitochondrial morphology due to infection, cells were stained with MitoTracker Deep Red and imaged using SIM (Supplementary Figure 5). We observed infection-induced changes to the mitochondrial network, in particular clustering around the gC-enriched perinuclear region. However, 2D imaging using SIM did not allow us to have a full overview of the mitochondrial network. Therefore, in order to gain a better picture of the mitochondrial network topography, we changed the imaging protocol and took z-stacks of infected cells that were fixed and immunostained for the mitochondria marker TOM20 using a widefield microscope. Stacks were then deconvolved and maximum intensity projected (Figure 5A). We observed a strong re-shaping of the mitochondria network morphology from a heterogeneous population of individual organelles (stage 1) to large interconnected tubular networks (stage 4). In particular, during stage 3, mitochondria appear to be densely concentrated around the gC-enriched perinuclear region. Stage 4 showed that these dense mitochondrial networks remain associated with fragmented gC-rich membranes, and these dense regions are connected by long tubular mitochondria.

**Figure 5:**
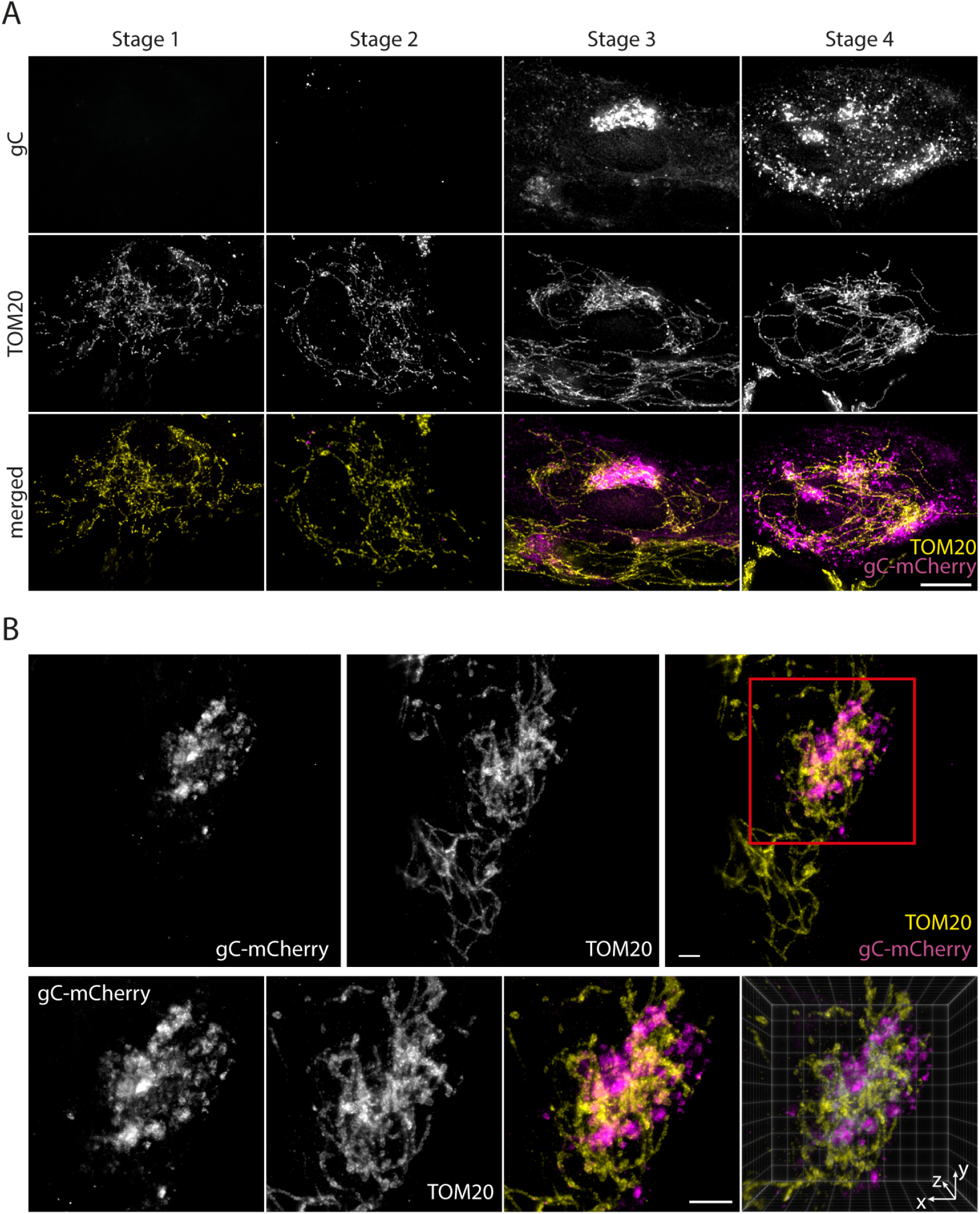
Interlacing of mitochondria and assembly compartments. **A** HFF cells were infected with eYFP-ICP0/gC-mCherry HSV-1, fixed at 3.5, 5.5, 7.5 and 9.5 hpi, and the mitochondria were labeled by staining TOM20 following standard immunofluorescence protocol. Z-stacks were taken at a widefield microscope. Displayed are maximum intensity projections after deconvolution showing exemplary cells for each stage of infection. From the top to bottom row: gC, mitochondria (TOM20) and the two channels merged (magenta: gC, yellow: mitochondria). Bottom row shows colour-encoded z-position of mitochondria. **B** Infected cells were fixed 9 hpi and immunostained for TOM20 to label mitochondria. mCherry signal was enhanced by use of a nanobooster. Samples were then expanded, and cells in stage 3 imaged using light sheet microscopy (Supplementary Figure 6). 3D structure of intertwined mitochondria and sites of secondary envelopment can be clearly observed. Upper row: whole cell, lower row: perinuclear region. Scale bar 10 μm.

However, the high concentration of mitochondria and gC-positive membranes prevented us from obtaining a clear overview of these convoluted structures using widefield microscopy. To address this, we employed the recently developed technique of expansion microscopy in combination with light sheet microscopy to resolve the 3D arrangement of mitochondria and gC-rich perinuclear compartment with much higher resolution. Expansion microscopy involves physical, isotropic swelling of fixed samples that can provide up to 64-fold increased sample volume, effectively providing super resolution imaging data using conventional fluorescence microscopes (Supplementary Video 2, Supplementary Video 3). Using this technique, we demonstrate that mitochondria and viral assembly compartments are intimately intertwined, with the mitochondria interlacing the gC-enriched membranes (Figure 5B).

Peroxisomes have also been reported to be important signaling platforms for antiviral innate immunity. We labeled peroxisomes in infected cells by immunostaining for catalase and analysed the dependence of peroxisome size on the stage of infection. However, no obvious change in peroxisome location was observed, except potentially during stage 4 where peroxisomes seem to be located around gC-enriched compartments (Figure 6). However, this seeming clustering could also be due to infection-induced cell rounding. In addition, we did not observe any significant change in the size of peroxisomes across all four stages of infection, with a consistent organelle median area of 0.20–0.22 μm.

**Figure 6:**
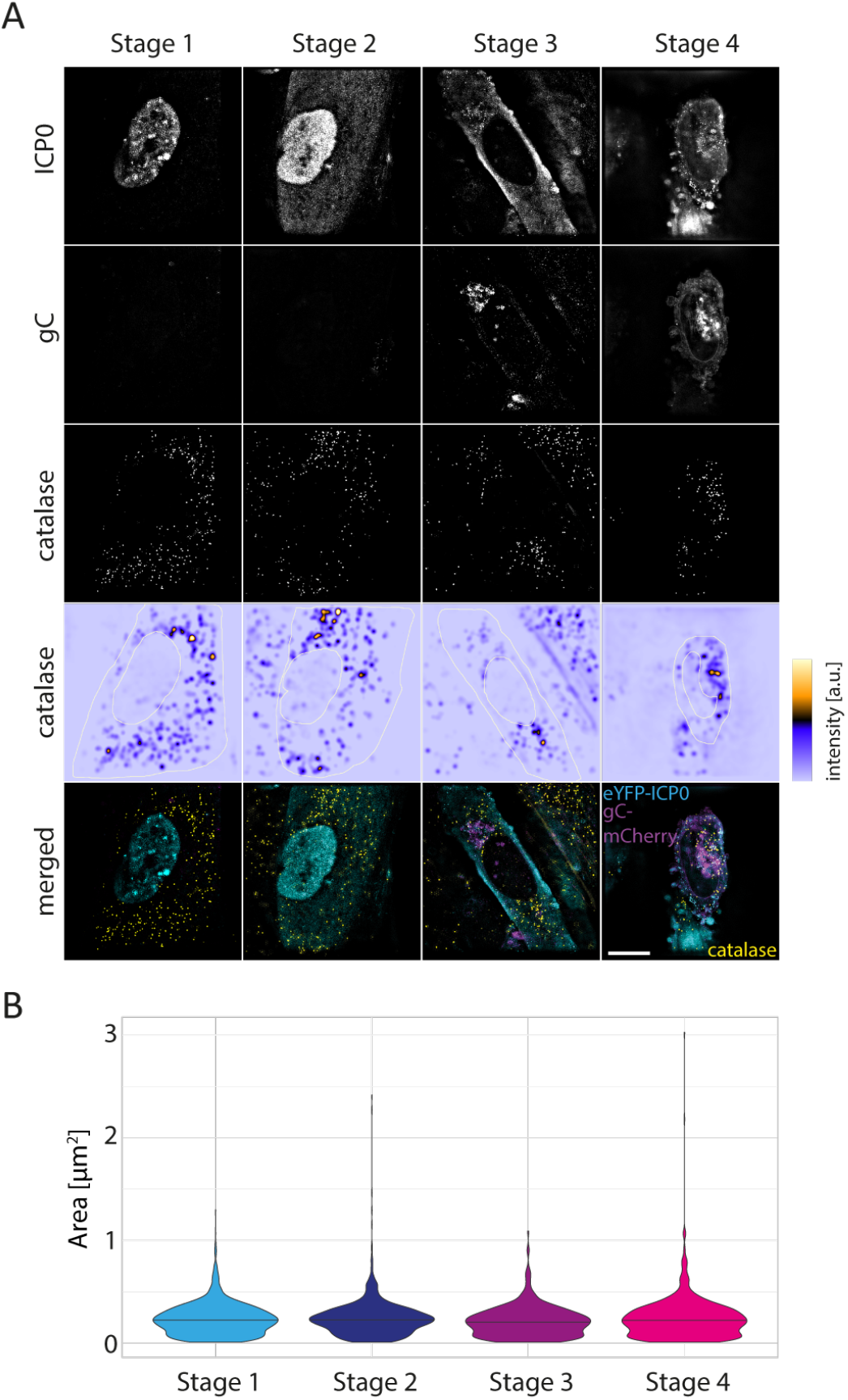
Spatial distribution of peroxisomes. **A** HFF cells were infected with eYFP-ICP0/gC-mCherry HSV-1, fixed at 3.5, 5.5, 7.5 and 9.5 hpi, and stained for peroxisomes with an antibody against catalase by use of a standard immunofluorescence protocol. No change of spatial distribution is seen. **B** Distribution of peroxisome size. The horizontal line in the violin plot indicates the median of the distribution. Scale bar 10 μm.

## Discussion

To understand the impact of viruses on host cell biology, as well as the mechanisms behind cytopathic effects and the wider pathogenesis of viral infection, it is vital to accurately define virus dependent changes to the subcellular architecture of cells. Herpesviruses are large and complex viruses that cause dramatic effects on cell morphology during their lytic replication cycles. However, there is a lack of understanding of when and how the architecture of a cell is changed during the course of herpesvirus infection, and in particular, how specific effects on morphology relate to phases of the replication cycle. We set out to address this by using a newly developed reporter virus that provides an accurate readout of infection stage and advanced high resolution microscopy analysis of multiple aspects of host cell morphology.

By engineering a virus with fluorescent tags fused to one of the earliest (ICP0) and one of the latest (gC) expressed viral proteins, we have created an intrinsic timestamp reporter of HSV-1 replication which allows for the accurate determination of the stage in viral replication at the single cell level. Observing the temporal and spatial expression patterns by use of high-resolution fluorescence imaging provides unambiguous visual descriptors to introduce a classification scheme, sorting each cell into one of four distinct stages of infection (stage 1 – stage 4). In the future, we envisage opportunities to further refine the classification of HSV-1 replication by including additional fluorescently tagged viral proteins with distinct spatial and kinetic expression profiles as well as through the use of machine learning to automate data categorisation (for reviews see Moen et al. [25] and von Chamier et al. [26]).

The benefits of our timestamping and classification method were demonstrated by studying the dynamics of viral replication at different MOIs. Our measured data fit well with the theoretically expected infection ratios in the cell population, but importantly highlight the degree of heterogeneity between individual cells within infected cell populations with co-occurrence of the different stages of infection at each time point. Without an intrinsic method to categorise the infection stage within each cell, such as our timestamp reporter virus method, attempting to relate any observed cellular changes to viral replication kinetics is fraught with difficulty. While using a high MOI such as 30 PFU/cell provides a seemingly more synchronous infection profile, a substantial increase in the fraction of cells in stage 4 at relatively early times post infection is the main effect, which reduces the opportunity to observe early stages of the infection process.

Using our timestamping methodology, we provide the first detailed overview of virus-induced changes to a broad range of host cell structures and organelles with a temporal correlation to the HSV-1 replication cycle. We investigated three classes of cellular structures involved in the HSV-1 life cycle: the cytoskeleton (microtubules and actin), secretory and endocytic compartments (the Golgi apparatus and early endosomes) and known antiviral signalling platforms (mitochondria and peroxisomes). The observed changes are summarised in Figure 7.

**Figure 7:**
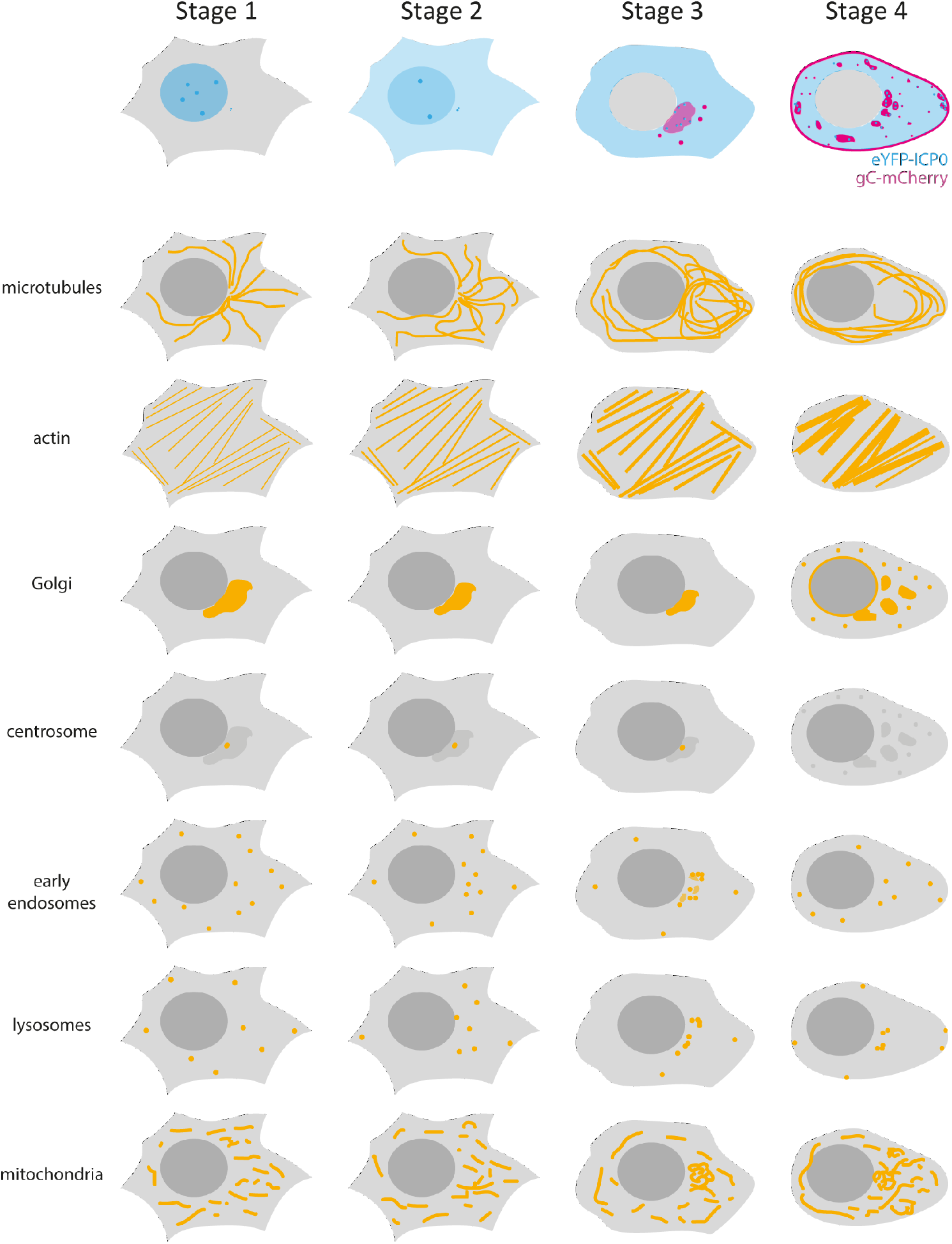
Overview of spatiotemporal morphological changes determined by use of the timestamping virus.

We discovered two main subsets of concerted morphological changes. First, and most obvious, are the changes that occur late in infection (stage 4). Almost all cellular structures and organelles that we monitored underwent profound rearrangement by this stage (>9.5 hpi). Due to experimental differences in cell line and experimental conditions such as hpi and MOI, it can be difficult to compare our data with previously stated morphological changes observed during HSV-1 infection. However, in an attempt to compare the structural changes at stage 4 with literature reports we selected studies which used high MOI and late times post infection, conditions for which cells in stage 4 should dominate the population. We found that our observations matched most descriptions of host cell rearrangements like redistribution and bundling of microtubules [27, 28] (13-16 hpi, MOI 3-5), loss of centrosome/MTOC [28] (13 hpi, MOI 5), migration of mitochondria towards perinuclear region [29] (6 hpi, MOI 5), and fragmentation of the Golgi apparatus [11, 27] (8-12 hpi, MOI 3-10) very well. An exception were peroxisomes, where we did not observe previously described morphological changes, possibly because they were previously identified at later times post infection (16 hpi, MOI 5; Jean Beltran et al. [30]).

Fragmentation of the Golgi complex and loss of the centrosome, such as that observed by stage 4 of infection, have also been described during apoptosis [31, 32]. Therefore, it seems plausible that during stage 4 morphological changes are caused by cytotoxic effects due to overloading the cell with late virus gene products and mature virions, rather than specific cell remodelling events by the virus to enhance replication. This correlates with the appearance of membrane blebbing and spikes that we observe during stage 4, which are also morphological characteristics of apoptosis [33].

Interestingly, we were able to detect an additional subset of concerted morphological changes that take place during the transition from stage 2 to stage 3. During this transition, the late viral genes are expressed which results in synthesis of the structural viral proteins necessary for virus assembly. Thus, we assume that the changes we observe are correlated to virus assembly. Central to the cellular rearrangement during the transition from stage 2 to stage 3 is the juxtanuclear region where the MTOC resides. Whereas microtubules seem to be pushed away from this part of the cell, early endosomes become clustered within this region alongside compaction of the Golgi complex, while mitochondria become intertwined with the juxtanuclear compartment enriched with viral glycoproteins. Previous studies speculated that mitochondria are accumulated within this region of the cell to supply energy for virus maturation, although no prominent activation of mitochondrial function could be observed, and a mechanism is still unknown [29]. The relocation of early endosomes and potential fusion with the Golgi compartment that we observed fit well with the model of viral assembly where secondary envelopment occurs at membranes originating from the endocytic pathway and the trans-Golgi network.

The role of microtubules in herpesvirus transport and egress is well established. However, the clearance of microtubules away from the juxtanuclear region around the centrosome/MTOC is puzzling but might be related to inhibition of MTOC function. ICP0 has been reported to be responsible for the rearrangement of microtubules during HSV-1 infection [20] and could play a key role in this process. Our imaging data supports the role of ICP0 for microtubule reorganisation during viral replication as we observe ICP0 localisation to the centrosome/MTOC from stage 1 onwards. Notably, the colocalisation between ICP0 and the MTOC was more apparent in live than in fixed cells which may explain why this has not been reported before.

In summary, we have developed a new technology which enables the direct, visual read-out of the stage of viral replication on the single cell level. One advantage of our method lies in its applicability in both live and fixed cells, and this methodology could prove useful for the study of other important viral pathogens where viral genes with different kinetic and spatial expression patterns can be effectively tagged with fluorescent reporters. In addition to linking morphological remodelling of the host cell to phases during the viral replication cycle, the tools we have developed can also be used to investigate the temporal correlation between HSV-1 replication and changes in, for example, cellular dynamics, mechanical properties and metabolic function. In the future, our approach can also be used to identify viral genes that are involved in the remodeling of the host cell through the introduction of mutations into the timestamping virus. This will therefore pave the way for gaining substantial new understanding on virus-host interaction based on high-resolution single-cell imaging.

## Materials and methods

### Antibodies and reagents

Primary antibodies anti-EEA1 (ab70521), anti-58K Golgi (ab27043), anti-pericentrin (ab28144) and anti-catalase (ab110292) were purchased from Abcam, anti-Tom20 (sc-17764) from Santa Cruz Biotechnology. All primary antibodies were monoclonal mouse antibodies. Polyclonal secondary anti-mouse antibodies conjugated to Atto647N (50185, Sigma Aldrich, Darmstadt, Germany) as well as isotype-specific secondary anti-mouse antibodies conjugated to Atto647N (610-156-041, Rockland Immunochemicals, Limerick, USA) and Alexa Fluor 568 (A-21124, Thermo Fisher Scientific, Waltham, USA) were used for detection. The nanoboosters - single-domain alpaca antibody fragments covalently coupled to fluorescent dyes — GFP-Booster Atto488 and RFP-Booster Atto594 (Chromotek, Martinsried, Germany) — were used to boost eYFP and mCherry fluorescence, respectively. Live cell stains SiR-tubulin, SiR-actin and SiR-lysosome were from Spirochrome (Stein am Rhein, Switzerland) and MitoTracker Deep Red FM was from Thermo Fisher Scientific (Waltham, USA).

### Cell lines and plasmids

Human foreskin fibroblast (HFF-hTERT) cells (American Type Culture Collection) were cultured with Dulbecco’s Modified Eagle’s Medium (DMEM, high glucose) supplemented with 10% fetal bovine serum, 1% GlutaMAX, 100 U/mL penicillin and 100 μg/mL streptomycin at 37°C and 5% CO_2_. Plasmids mIFP12-Rab4a-7 (#56261) and mIFP-Golgi-7 (#56221) were purchased from Addgene.

### Cloning and generation of mIFP-expressing stable cell lines

The mIFP-tagged constructs were amplified by PCR using primers containing flanking Gateway attB sequences and cloned into the Gateway donor vector pDONR223. Constructs were subsequently cloned into lentiviral destination vector pHAGE-pSFFV using the Gateway system (Thermo Scientific). Lentiviral particles were generated by transfection of HEK293T cells with the pHAGE-pSFFV vectors plus four helper plasmids (VSVG, TAT1B, MGPM2, CMV-Rev1B), using TransIT-293 transfection reagent (Mirus) according to the manufacturer’s recommendations. Lentiviral supernatants were harvested 48 h after transfection, cell debris was removed with a 0.22 μm filter, and HFF-hTERT cells were transduced for 48 h then subjected to puromycin selection to generate stable cell populations expressing mIFP-tagged markers.

### Recombinant viruses

Fluorescently labelled, recombinant eYFP-ICP0/gC-mCherry virus was used as a visual reporter for the stage of infection. The ‘timestamp’ recombinant reporter virus was constructed using the bacterial artificial chromosome (BAC)-cloned KOS strain of HSV-1 [34], and the two-step Red recombination technique [35]. Three rounds of two-step Red recombination were conducted to firstly insert the eYFP(A206K) coding sequence in place of the initiating ATG of both copies of the RL2 gene (ICP0) followed by the insertion of the mCherry coding sequence immediately before the stop codon of UL44 (gC). Infectious virus was reconstituted by transfection of Vero cells with BAC DNA together with a Cre recombinase expression plasmid to excise the BAC-cassette. All virus stocks were grown in Vero cells and infectious titres determined by plaque assay on HFF-hTERT and Vero cells.

### Infection assay

For infection assays, HFF cells were plated in 8-well LabTek coverglass chambers at 25,000 cells per well or onto 13 mm diameter round coverslips in 4-well plates at 60,000 cells per well 12 hours prior to infection. The next day, cells were infected with recombinant virus at 3 PFU per cell if not indicated otherwise. After one hour of incubation at 37°C and 5% CO_2_, medium was exchanged. Depending on the experiment, samples were transferred onto the respective microscope equipped with appropriate environmental control for live cell imaging after 3hpi or fixed, permeabilised and labelled with antibodies at different hpi as indicated and according to procedures described below.

### Immunostaining

Cells were immunostained using a standard immunofluorescence protocol. For fixation, cells were incubated in PBS with 4% methanol-free formaldehyde (28906, Thermo Fisher Scientific, Waltham, USA) for 10 min at room temperature. After permeabilisation in PBS with 0.2% (v/v) TritonX-100 for 5 min and blocking with 10% goat serum in PBS for 45 min, samples were incubated at room temperature for one hour with primary antibody (1:200 dilution) in PBS solution containing 3% bovine serum albumin (BSA). After washing three times with PBS, cells were incubated with Atto647N-labelled secondary antibody (1:400 dilution in PBS with 3% BSA) for 30 min at room temperature in the dark. After washing three times with PBS, cells were post-fixed with formaldehyde as described before.

Samples that were expanded as described below were first fixed with 4% methanol-free formaldehyde (28906, Thermo Fisher Scientific, Waltham, USA) and 0.1% glutaraldehyde (G7651, Sigma Aldrich, Darmstadt, Germany) in PBS for 10 min, and permeabilised by incubation with a Tween 20 solution (0.5% v/v in PBS) for 15 minutes at room temperature. After one wash in PBS, cells were incubated with blocking buffer (10% goat serum and 0.05% Tween 20 in PBS) for 30 minutes at room temperature. Samples were then incubated with the primary antibody (1:200 dilution) in blocking buffer at 4°C overnight, washed three times with PBS over 15 minutes, and incubated with the conjugated secondary antibody (1:400 dilution) in blocking buffer for one hour at room temperature in the dark. After washing three times with PBS over 15 minutes, a nanobooster was used to enhance fluorescence of mCherry-tags with a synthetic dye which is brighter and more resistant to the expansion protocol than the fluorescent protein tags. Cells were incubated with nanoboosters for eYFP and mCherry diluted 1:200 in PBS containing 4% BSA and 0.05% Tween 20 for 1 hour at room temperature in the dark. Samples were washed 3 times with PBS over 15 minutes and expanded the next day.

### Gelation, digestion and expansion of HSV-1 infected cells

Infected, fixed and immunostained cells were expanded following the expansion protocol described by Chozinski et al. [36] which is compatible with conventional synthetic dyes and autofluorescent protein tags. Fixed cell samples on 13 mm diameter round coverglass were incubated in monomer solution (2 M NaCl, 2.5% w/w acrylamide, 0.15% w/w N,N’-methylenebisacrylamide, 8.625% w/w sodium acrylate in PBS) for 1 minute at room temperature prior to gelation.

Concentrated stocks of ammonium persulfate (APS) and tetramethylethylenediamine (TEMED) at 10% (w/w) in water were diluted in monomer solution to concentrations of 0.2% (w/w) for gelation, with the initiator (APS) added last. The gelation solution (70 μl) was placed in a 1 mm deep, 1 cm diameter Teflon well and the coverglass was placed on top of the solution with cells face down. After 30 min at room temperature, gelation was complete. The coverglass and gel were removed with tweezers and placed in digestion buffer (1 ×TAE buffer, 0.5% Triton X-100, 0.8 M guanidine HCl) containing 8 units/mL proteinase K (17916, Thermo Fisher Scientific, Waltham, USA) freshly added. Gels were digested at 37°C for a maximum of 30 min. The gels were removed from digestion buffer and placed in 50 mL DI water to expand. Water was exchanged every 30 min until expansion was complete (typically 3–4 exchanges).

### Widefield microscopy

Time-lapse imaging of live HSV-1 infected cells was carried out at a custom-built widefield microscope. Frame (IX83, Olympus, Tokyo, Japan), stage (Prior, Fulbourn, UK), Z drift compensator (IX3-ZDC2, Olympus, Tokyo, Japan), plasma lightsource (HPLS343, Thorlabs, Newton, USA), and camera (Clara interline CCD camera, Andor, Belfast, UK) were controlled by Micro-manager [37]. Respective filter cubes for YFP (excitation 500 nm, dichroic mirror 515 nm, emission 535 nm), mCherry (excitation 560 nm, dichroic mirror 585 nm, emission 630 nm) and DAPI (excitation 350 nm, dichroic mirror 353 nm, emission 460 nm) were used. Images were acquired with an Olympus PlanApoU 60×/1.42 oil objective lens at 20–25 random positions for each sample. For 12 hours, images were recorded every 20 min starting at 3.5 hpi. During the experiment, cells were heated at 37°C and supplied with 5% CO_2_ via a stage top incubator and temperature, gas and humidity controllers (Okolab, Pozzuoli, Italy).

Volumes of fixed cells immunostained for TOM20 were acquired as z-stacks with a step size of 0.3 μm over a range of 9 μm. Data were deconvolved and maximum intensity projected as described below.

### Structured illumination microscopy (SIM)

Structured illumination images were collected on a custom-built structured illumination microscopy (SIM) setup, modified from one which has been described before in detail [38]. A 60×/1.2NA water immersion lens (UPLSAPO60XW, Olympus, Tokyo, Japan) was used to focus the structured illumination pattern onto the sample and captured the samples’ fluorescence emission light which was detected with a sCMOS camera (C11440, Hamamatsu, Hamamatsu-City, Japan). Laser excitation wavelengths used were 488 nm (iBEAM–SMART-488, Toptica, Graefelfing, Germany), 561 nm (OBIS561, Coherent, Santa Clara, USA), and 640 nm (MLD640, Cobolt, Solna, Sweden). Respective emission filters were BA510-550 (Olympus, Tokyo, Japan), BrightLineFF01-600/37, and BrightLineFF01-676/29 (Semrock, NewYork, USA). Imaging was done in fixed or live cells, as indicated. During the live cell experiment, cells were heated at 37°C and supplied with 5% CO_2_ via a stage top incubator and temperature, gas and humidity controllers (Okolab, Pozzuoli, Italy). Images were acquired using HCImage (Hamamatsu, Hamamatsu-City, Japan) and a custom-built control software based on Labview (National Instruments, Newbury, UK). Nine raw images were collected at each plane and each color. Raw SIM images were reconstructed with fair-SIM [39] using a customised Fiji macro including correction for the lateral shift between colours, measured using 100 nm beads (TetraSpeck Microspheres, Thermo Fisher Scientific, Waltham, USA). Out-of-focus light was suppressed by intermixing separated bands as suggested by O’Holleran & Shaw [40].

### Light sheet microscopy

Expanded samples were imaged at a custom-built inverted selective plane illumination microscope (iSPIM). Parts were purchased from Applied Scientific Instrumentation (ASI, Eugene, USA) including controller (TG8_BASIC), scanner unit (MM-SCAN_1.2), right-angle objective mounting (SPIM-K2), stage (MS-2K-SPIM) with motorised Z support (100 mm travel range, Dual-LS-100-FTP) and filter wheel (FW-1000-8). All components were controlled by Micro-Manager by means of the diSPIM plugin. The setup was equipped with a 0.3 NA excitation objective (Nikon 10×, 3.5 mm working distance) and a higher, 0.9 NA detection objective (Zeiss, W Plan-Apochromat 63×, 2.1 mm working distance) to increase spatial resolution and fluorescence signal collection. Lasers (OBIS488-150 LS, OBIS561-150 LS and OBIS647-120 LX, Coherent, Santa Clara, USA) were fiber-coupled into the scanner unit. An sCMOS camera (ORCA-Flash 4.0, Hamamatsu, Hamamatsu-City, Japan) was used to detect fluorescence. Respective emission filters were BrightLineFF01-540/50, BrightLineFF01-624/40 and BrightLineFF0-680/42 (Semrock, NewYork, USA). Gels containing expanded samples were cut into small strips and mounted onto 24 mm × 50mm rectangular coverslips, with expanded cells facing upwards, using Loctite super glue (Henkel, Duesseldorf, Germany). The sample was then placed into an imaging chamber (ASI, I-3078-2450) which was filled with water. We recorded 150 planes per volume and colour channel, spacing planes every 0.5 μm. Raw data were deskewed using a custom ImageJ plugin.

### Cell counting and assignment of stage of infection

Cells were seeded onto 13 mm diameter round coverslips as described above, and infected the next day with eYFP-ICP0/gC-mCherry virus at MOI 0.3, 3 and 30, respectively. Samples were fixed every hour after infection, starting at 3 hpi until 8 hpi. In order to increase fluorescent signal of eYFP- and mCherry-tags of the reporter virus, cells were immunostained with nanoboosters. Samples were then mounted in mounting medium containing DAPI (VECTASHIELD HardSet Mounting Medium, Vectorlabs, Burlingame, USA) according to standard manufacturer’s protocol. For each condition, 20–25 fields-of-view (FOVs) were acquired at the widefield microscope as described above with 5–20 cells per FOV. Nuclei were counted to yield the total number of cells and each individual cell was then classified as non-infected or assigned a stage of infection according to the classification scheme. For each condition, between 135–300 cells were counted.

### Measurement of organelle area

The area occupied by the Golgi complex and peroxisomes was determined from reconstructed SIM images using ImageJ. For the Golgi complex, the outline of the compartment was traced manually, whereas for peroxisomes binary images were created by use of Otsu thresholding. The area of the compartments was then determined using the ‘Analyze Particles’ function. Violin plots were generated by use of the web app PlotsOfDifferences [41].

### Density maps

SIM images of vesicular structures were Gaussian filtered (10 pixel) in order to smooth edges of fluorescent structures using ImageJ. For highlighting regions with a high density of vesicular structures, the lookup table ICA was chosen.

### Deconvolution

Z-stacks taken at the widefield microscope and deskewed lightsheet microscopy data were deconvolved using a custom GPU-accelerated chunked deconvolution routine written in MATLAB. 250 iterations of the unregularised Richardson-Lucy algorithm were used on an initial estimate set to the mean value of the raw data, following a background subtraction to correct for camera offset. Deconvolved data were maximum intensity projected in ImageJ, using colour to indicate depth as indicated.

## Author Contributions

K.M.S. prepared samples, performed experiments as well as data analysis and wrote the manuscript. J.D.M. designed and built imaging hardware, and developed data processing and analysis software. T.K.S. made viral stocks and determined titers. L.M. prepared expanded samples. V.C. generated the eYFP-ICP0/gC-mCherry HSV-1. V.C. and C.M.C. made stable cell lines. C.F.K. and C.M.C. supervised the project.

## Supplementary Information

**Supplementary Figure 1:**
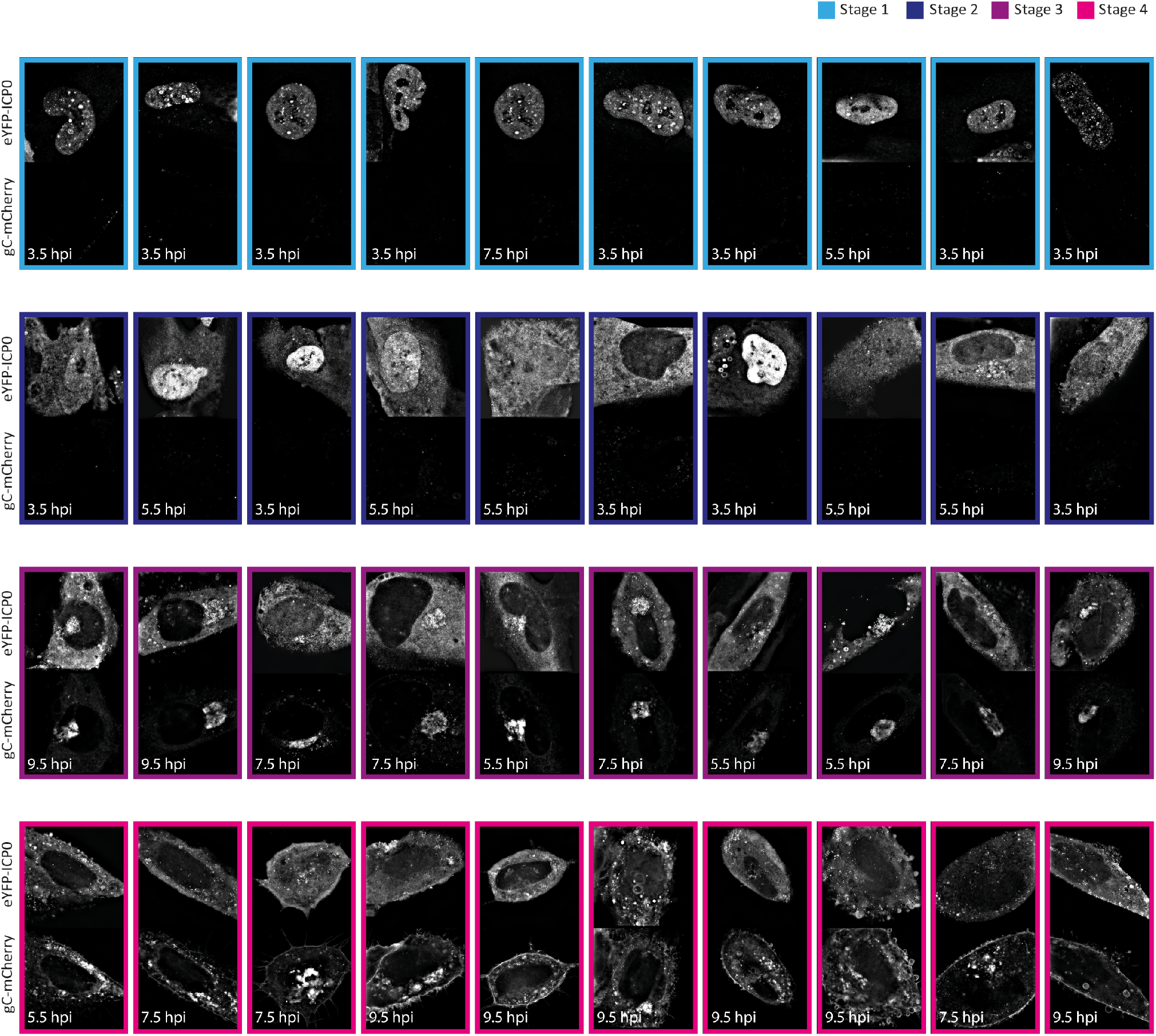
Variation in the stage of infection at the individual cell level at different hours post infection. HFF cells were infected with eYFP-ICP0/gC-mCherry HSV-1 and fixed at 3.5, 5.5, 7.5 and 9.5 hpi. Visual features determined by eYFP-ICP0 and gC-mCherry appearance and distribution were used to classify each individual cell. Despite different cell shapes, features are mostly uniform for each stage of infection. Exception is stage 2 which is characterised by the transition of ICP0 from the nucleus into the cytoplasm. Individual cells captured during the transition can appear with ICP0 being present still predominantly in the nucleus, with equal amounts in the nucleus and the cytoplasm, or predominantly in the cytoplasm in different cells.

**Supplementary Figure 2:**
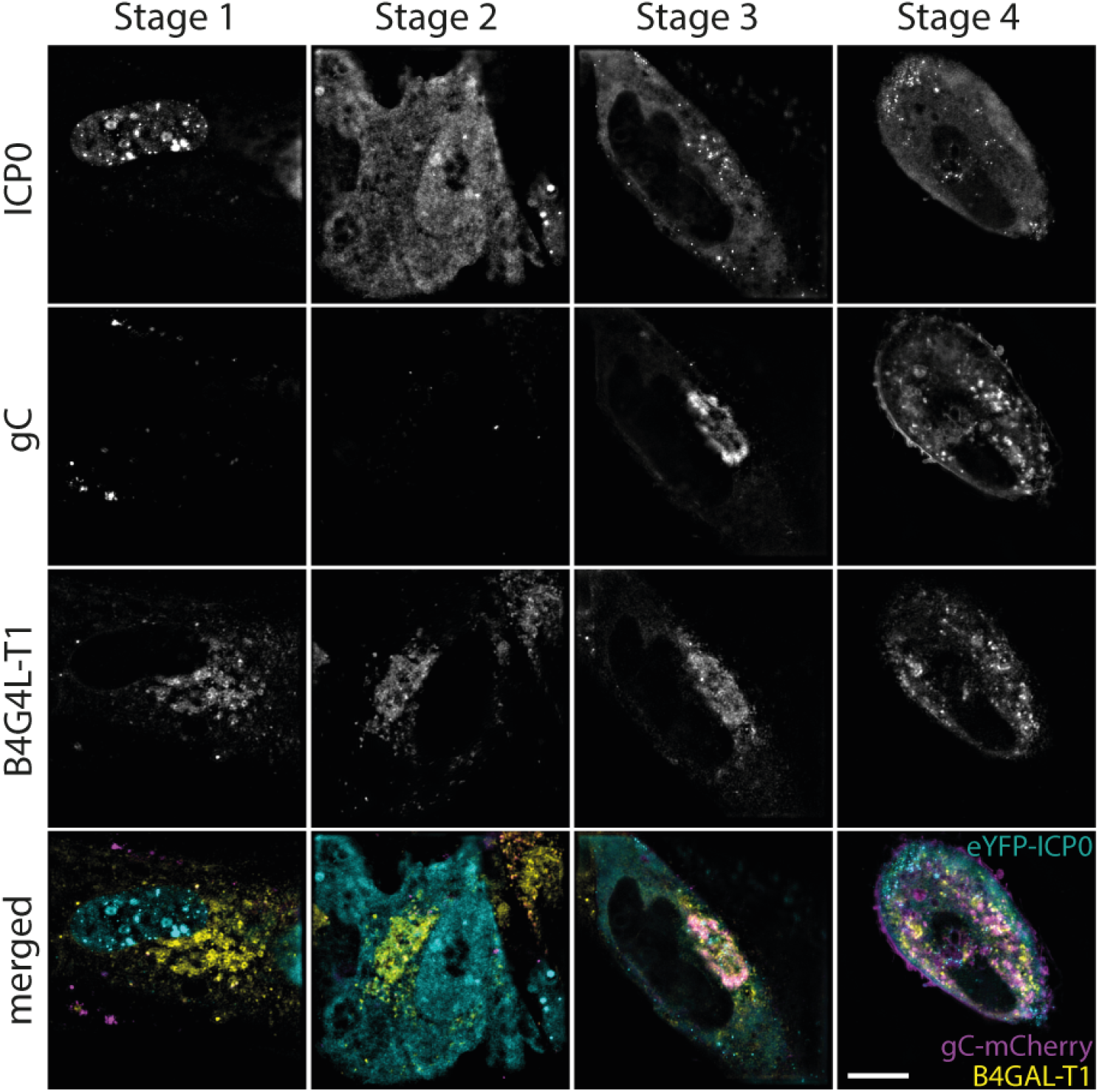
Morphology changes and fragmentation of the Golgi complex during HSV-1 replication. HFF cells stably expressing mIFP-B4GAL-T1 were infected with eYFP-ICP0/gC-mCherry virus and fixed at 3.5, 5.5, 7.5 and 9.5 hpi. Scale bar 10 μm.

**Supplementary Figure 3:**
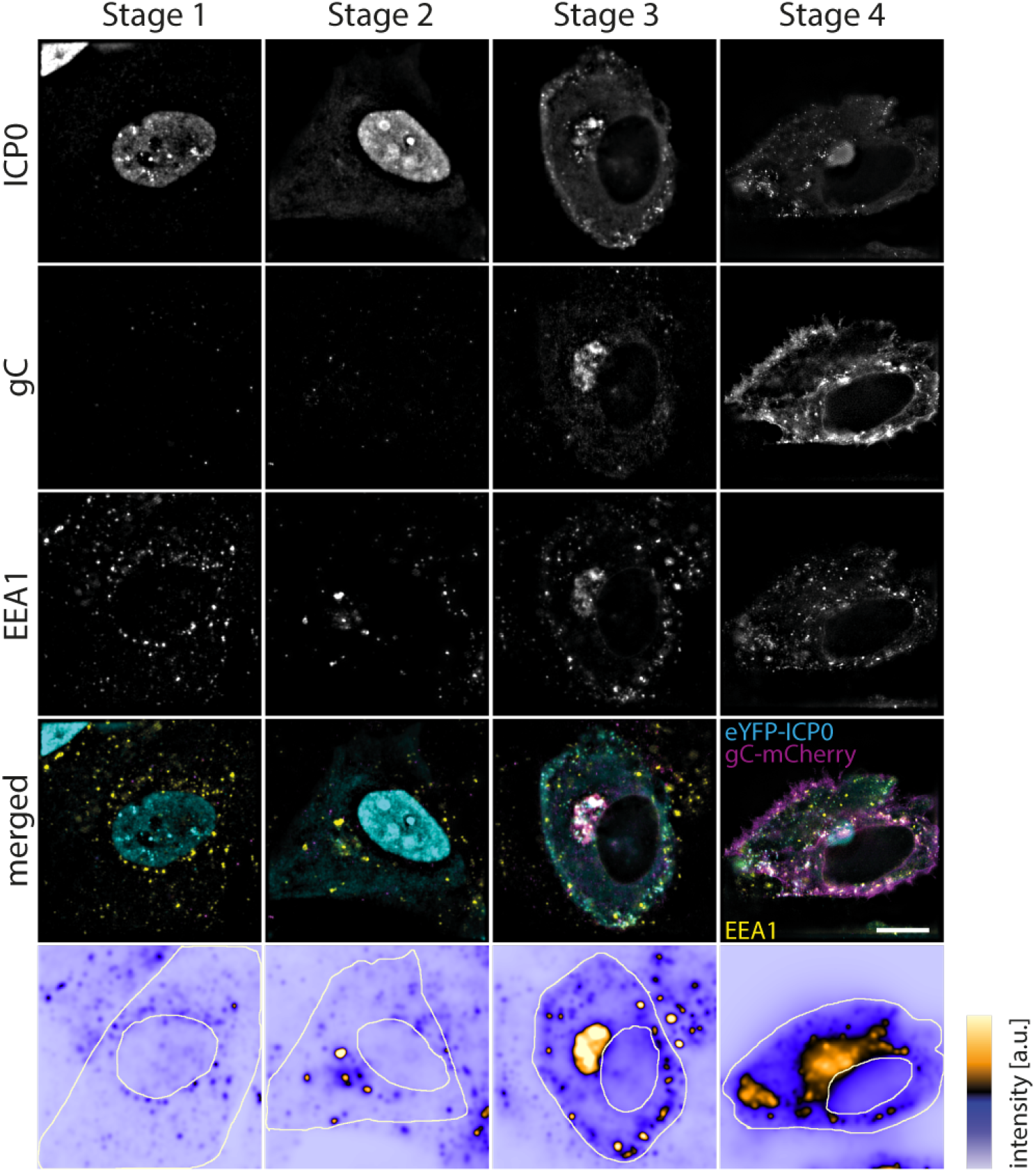
Spatial distribution of early endosomes. HFF cells were infected with eYFP-ICP0/gC-mCherry HSV-1, fixed at 3.5, 5.5, 7.5 and 9.5 hpi, and stained for early endosomes with an antibody against EEA1 by use of a standard immunofluorescence protocol. Early endosomes accumulate at the perinuclear region during stage 2 and 3, and re-distribute in the cytoplasm during stage 4 due to fragmentation of perinuclear compartment. Scale bar 10 μm.

**Supplementary Figure 4:**
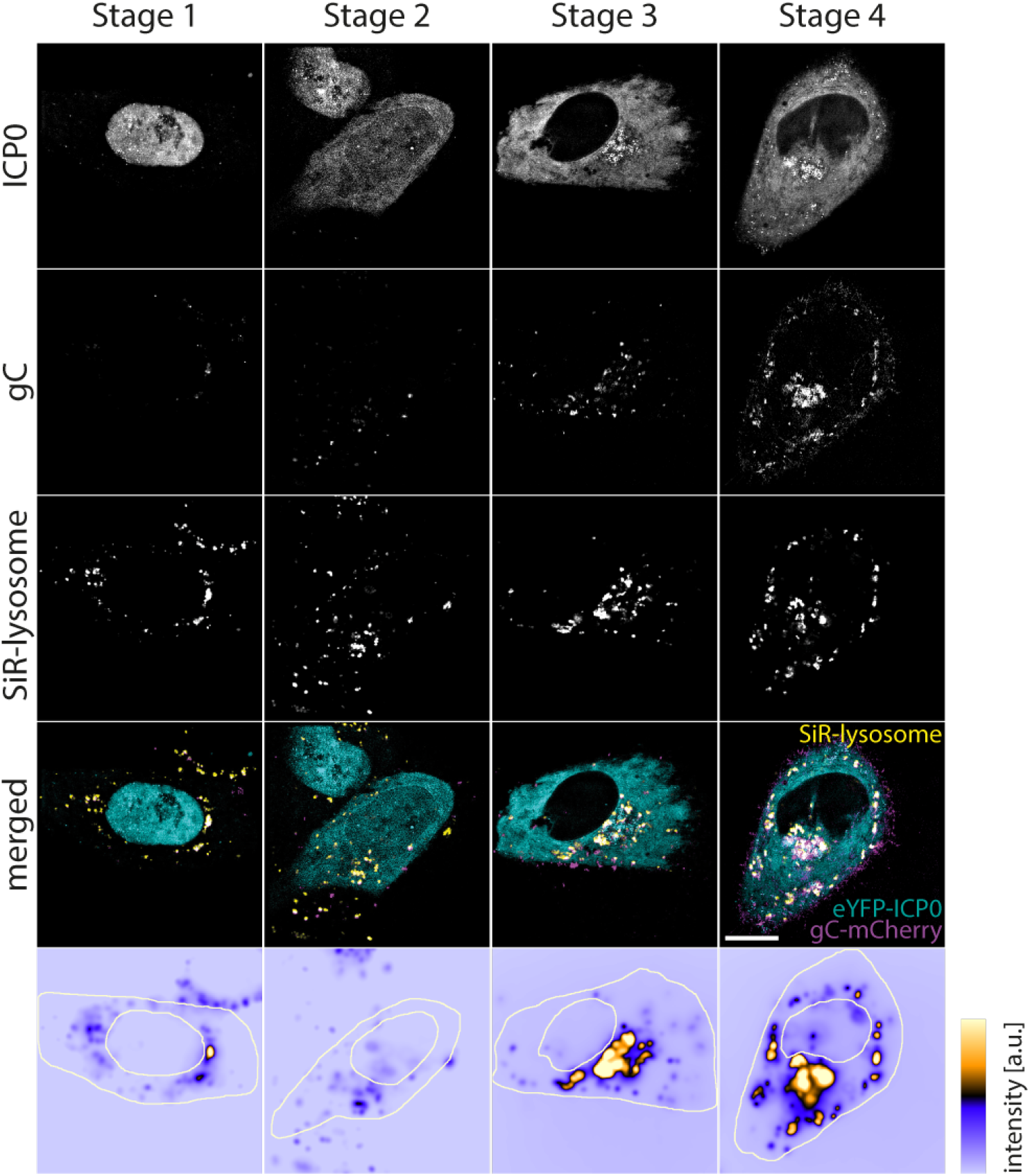
Lysosome distribution during all stages of infection. HFF cellswere infectedwith eYFP-ICP0/gC-mCherry HSV-1, stained with SiR-lysosome and imaged at 3.5, 5.5, 7.5 and 9.5 hpi. Lysosomes are attracted to compartments rich in gC. Scale bar 10 μm.

**Supplementary Figure 5:**
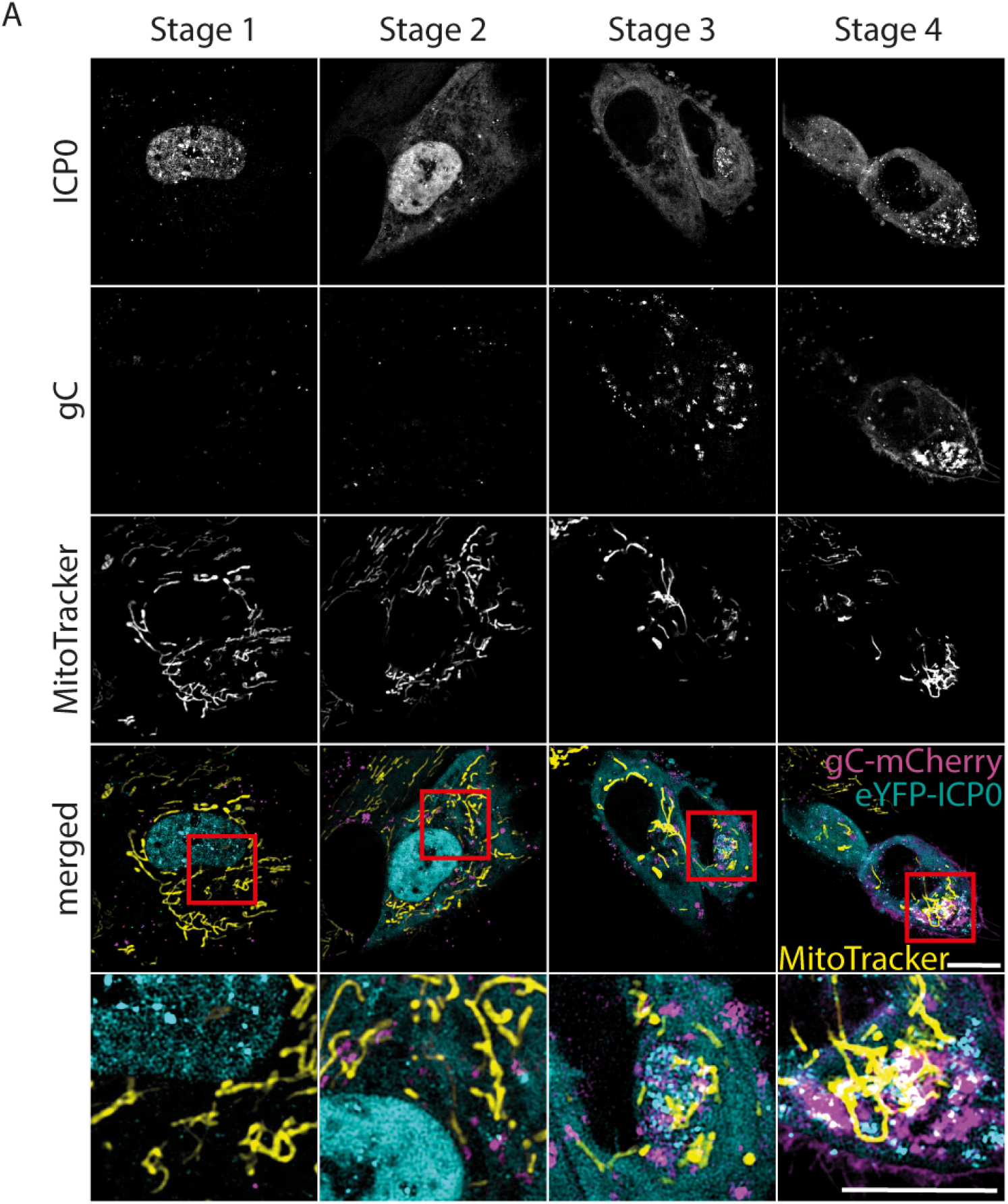
Mitochondria re-location towards gC-rich compartments. HFF cells were infected with eYFP-ICP0/gC-mCherry HSV-1, stained with MitoTracker Deep Red and imaged at 3.5, 5.5, 7.5 and 9.5 hpi. Scale bar 10 μm.

**Supplementary Figure 6:**
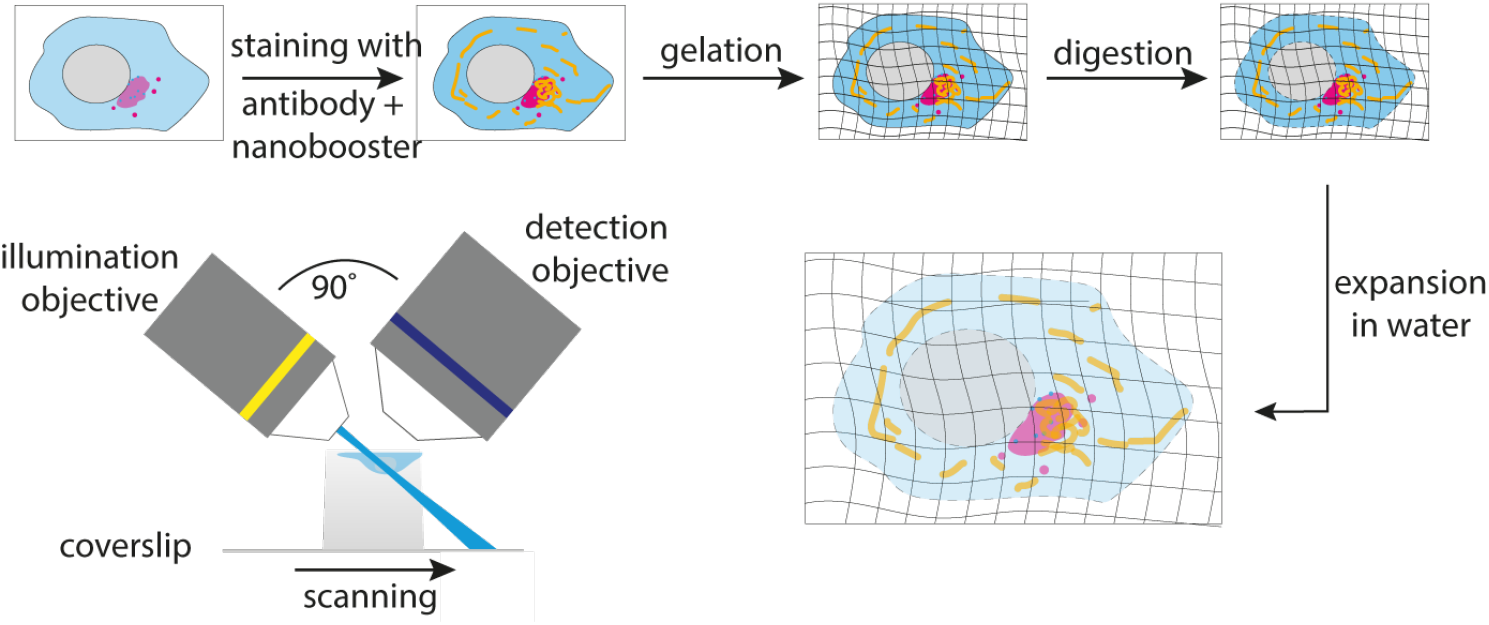
Schematic illustration of expansion microscopy and light sheet microscopy. Infected cells were fixed, and structures of interest were stained using a standard immunofluorescence protocol. A nanobooster specific for mCherry was used to enhance the fluorescence signal from the viral protein gC. Expanded samples were cut into thin strips and mounted onto coverslips with superglue for imaging on a custom-built, inverted light sheet microscope.

**Supplementary Video 1:**
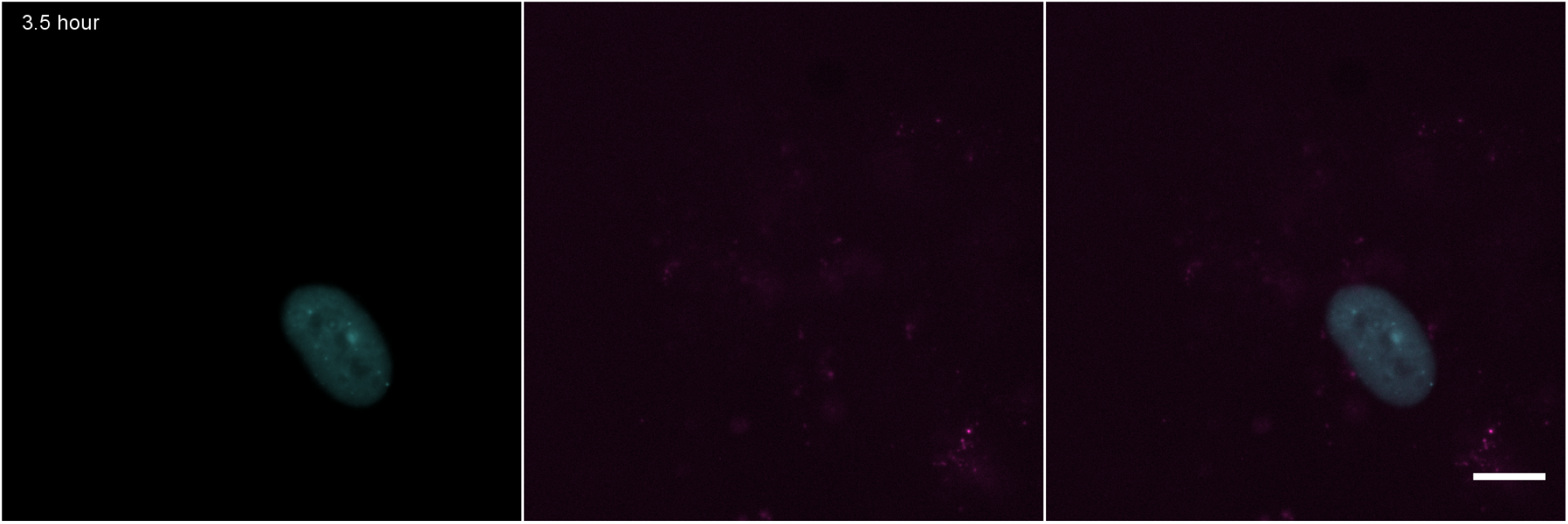
Time-lapse imaging of HFF cells infected with eYFP-ICP0/gC-mCherry HSV-1. Imaging was started 3.5 hpi, and images were recorded for 12 hours every 20 min. The transition of eYFP-ICP0 (cyan) from the nucleus into the cytoplasm as well as the expression of gC-mCherry (magenta) late in infection can be clearly observed, confirming the functionality of our reporter virus. Scale bar 10 μm.

**Supplementary Video 2:**
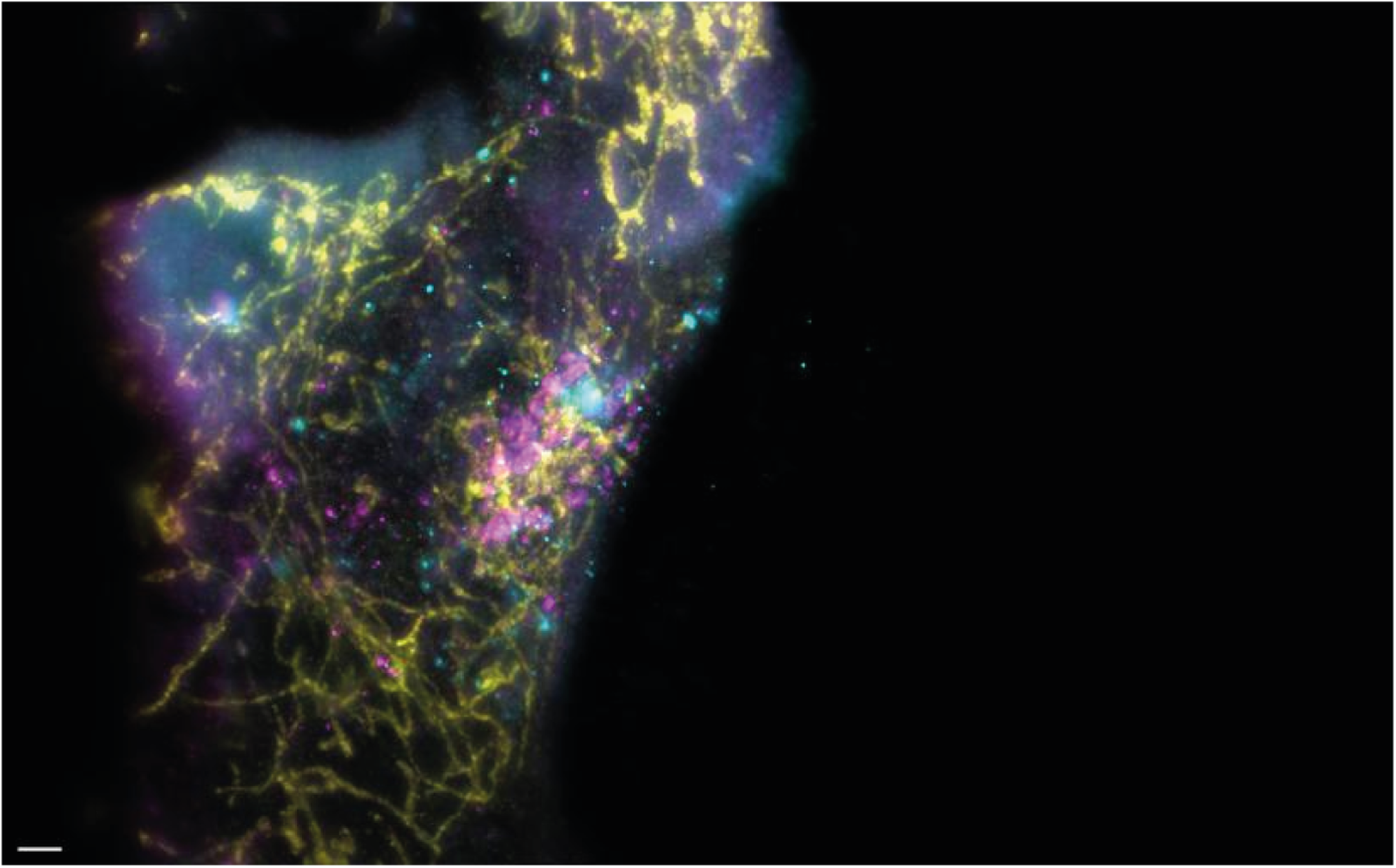
Interlacing of mitochondria with compartment of secondary envelopment at stage 3 of infection. Infected cells were fixed 7.5 hpi, and immunostained for TOM20. Fluorescence of eYFP and mCherry was enhanced by use of nanoboosters. Samples were expanded 4 ×, and imaged at a custom-built light sheet microscope. The video corresponds to Figure 5B in the main text. Scale bar 10 μm. Cyan: ICP0, magenta: gC, yellow: TOM20.

**Supplementary Video 3:**
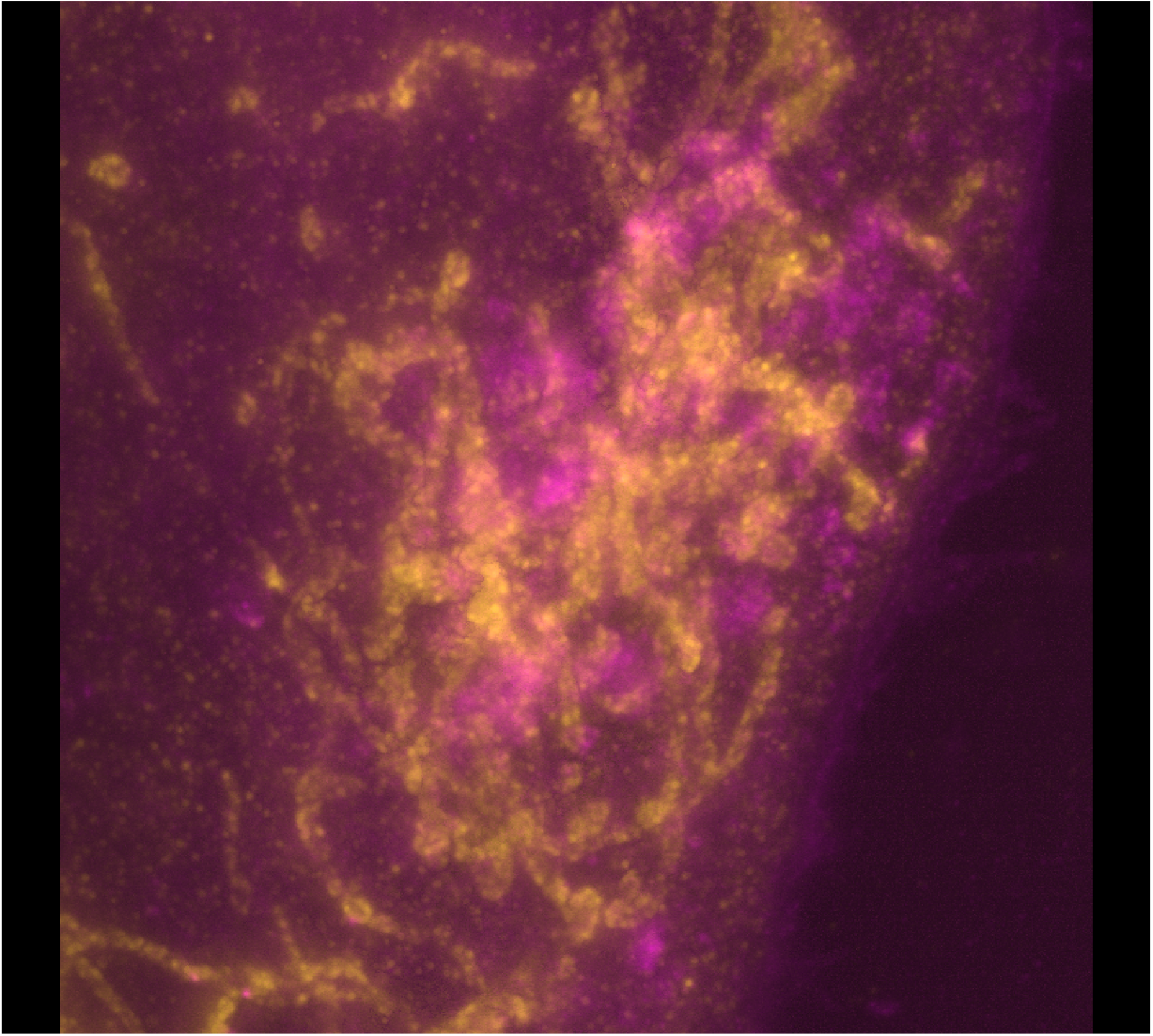
Enlarged view of Supplementary Video 2. Only gC (magenta) and TOM20 (yellow) are shown to highlight the interlacing of the mitochondria with the gC-enriched compartment. The video corresponds to Figure 5B (magnified section in bottom row) in the main text.

